# T32 training is associated with increased likelihood of obtaining an academic research faculty position: a cross-sectional study

**DOI:** 10.1101/2022.04.27.489715

**Authors:** Adrienne L. Mueller, Addie Schnirel, Sofie Kleppner, Philip Tsao, Nicholas J. Leeper

## Abstract

**Background:** The main mission of NIH-sponsored institutional training programs such as the T32 is to provide strong research and career training for early career scientists, while preparing those individuals to become leaders working to meet the health-related research needs of the nation. One of the main avenues to pursue health-related research is becoming research faculty at an academic institution. It is therefore important to know whether these programs are succeeding in this mission, or, if barriers exist that prevent trainees from pursuing these careers.

**Methods:** Our institution currently trains ~ 2400 post-doctoral scholars per year, approximately 5% of whom are enrolled in one of our 33 T32 programs. In this study, we 1) compare the professional outcomes of T32 trainees with non-T32 trainees at our institution, and 2) survey past and current T32 trainees in a subset of high-performing cardiovascular programs about the barriers and enablers they experienced to pursuing research-oriented careers.

**Results:** Former T32 trainees are significantly more likely to attain appointments as primarily research faculty members, compared to other trainees. Trainees report a perceived lack of stability, the paucity of open positions, and the ‘publish or perish’ competitive mentality of academia as their top reasons for abandoning careers in academia. However, they were still more likely to choose research over clinical careers after participating in a dedicated T32 program.

**Conclusions:** Our results support the conclusion that structured training programs strengthen the pipeline of young scientists pursuing careers in academic research, including those from underrepresented backgrounds. However, T32 postdoctoral researchers are held back from pursuing academic careers by a perceived lack of stability and high competition for faculty positions.

**Funding:** This research received no specific grant from any funding agency in the public, commercial, or not-for-profit sectors.

## Introduction

In 2018, the White House released the STEM Education Strategic Plan: a federal strategy for providing Americans with persistent, high-quality access to STEM education and to position the US as a global leader in STEM professions. A key component of this initiative is the National Institutes of Health’s (NIH) continued investment in formal training programs for early career researchers. While the NIH offers a range of educational awards, the Ruth L. Kirschstein Institutional National Research Service Award (also known as a T32 award) is considered the backbone of its training programs. The main goal of T32 programs is to provide strong research and career training for early career scientists, who will then continue on as leaders in STEM research, particularly in academic positions as research faculty. T32 programs prepare individuals who are committed to a research career to transition to their next career stage. These programs not only provide salary support for post-doctoral trainees, but also typically include a wide range of structured career development programming, including workshops, discussions and Individual Development Plans, to enable trainees’ successful transition to the next stage of their careers. The programs support dedicated research training time in the labs of faculty mentors, but also ensure that all trainees are well-versed in the fundamentals of practicing sound science mandating training in rigorous and reproducible research methods and the responsible conduct of research. It is therefore important to know whether these investments are succeeding in this goal, and what the factors are that contribute to, or inhibit, that success.

Our Tier 1 research institution currently hosts 33 distinct training programs spanning all aspects of human health and disease, including three that are specifically related to cardiovascular research. These programs support approximately 5% of our total post-doctoral trainee population, the remainder of whom do not necessarily participate in the structured programming offered via the T32 mechanisms. In this study, we 1) compare the career outcomes of T32 trainees with non-T32 trainees at our institution, and 2) summarize the surveyed reports of past and current trainees from a subset of high-performing T32 programs about the barriers and enablers they have experienced to pursuing careers in academia.

## Results

Based on the data from our institution’s Office of Postdoctoral Affairs dataset, approximately 40% of postdocs attain faculty appointments after their training at our institution (Figure 1A). There is no significant difference in the proportion of trainees who become faculty compared to non-faculty between T32 Postdocs and Non-T32 Postdocs (p = 0.42, d.f. = 1). Conversely, former-T32 trainees are significantly more likely to attain appointments as primarily research faculty members, compared to other trainees (Figure 1B) (p = 0.002, d.f. = 1). Though limited by a small sample size, we also observed that compared to those that did not participate in a T32, a numerically higher proportion of postdocs who self-identified as underrepresented minorities went on to obtain a primarily research faculty position if they had participated in a T32. Of the 205 URM postdocs, 31.3% of the T32 trainees achieved such positions, compared to only 16.9% of URM trainees who participated in traditional post-doctoral fellowships. This result was not significant at a threshold of 0.05 (p = 0.16 with Bonferroni correction.)

**Figure 1.**
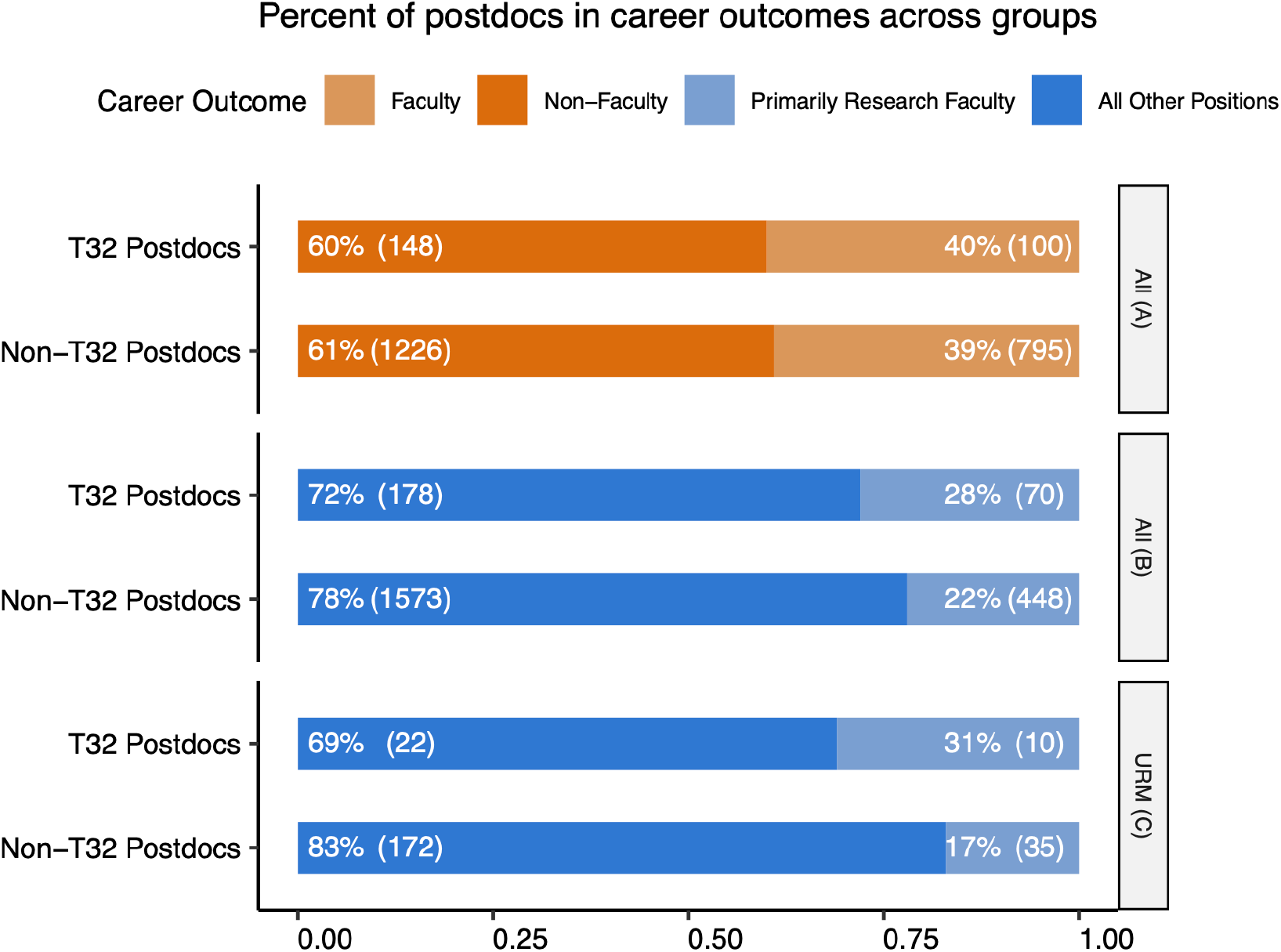
The proportion of trainees identified as holding either faculty or non-faculty positions for two populations: T32 Postdocs and Non-T32 Postdocs. B: The proportion of trainees who identified as holding either primarily research faculty positions or any other position for T32 Postdocs and Non-T32 Postdocs. C: Among trainees who identified as underrepresented minorities, the proportion trainees who identified as holding either primarily research faculty positions or any other position for two populations: trainees with T32 Postdocs or Non-T32 Postdocs. Data in panels A, B and C are from the OPA dataset. See accompanying deidentified source data: “OPA Data.”

Compared to the pooled outcomes from the non-cardiovascular T32 programs, trainees from the cardiovascular programs had a higher likelihood of attaining a primarily research faculty position: 34% compared to 26.7%, though the p-value was not significant at the 0.05 threshold (p = 0.3, d.f. = 1). With regards to the outcomes of trainees specifically in the three cardiovascular T32 programs, we find that trainees have a similar distribution of outcomes, regardless of graduate degree type (Figure 2). There was no significant difference in the distribution of outcomes across training background (p = 0.29, d.f. = 6). Notably, there was a low incidence of trainees entering clinical practice, regardless of MD, MD/PhD or PhD background. There was also no significant difference in the distribution of outcomes across cardiovascular T32 programs (p = 0.84, d.f. = 6, data not shown).

**Figure 2.**
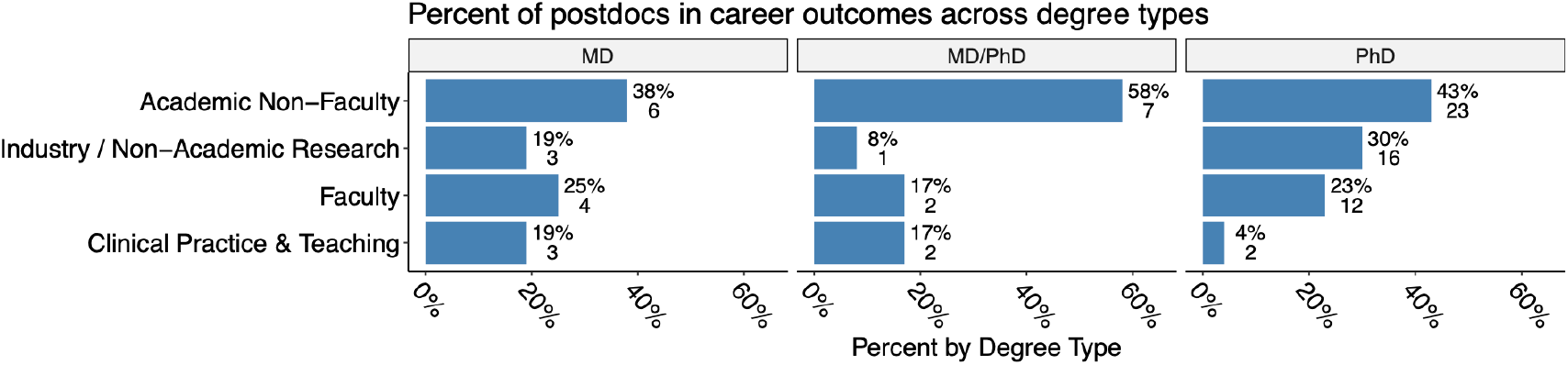
Distribution of T32 program trainees across different career sectors, categorized by their previous training background (MD, MD/PhD, PhD). Data from CV T32 Data. See accompanying deidentified source data: “CV T32 Program Data.”

We next surveyed trainees in the three cardiovascular T32 programs regarding the primary barriers and enablers to pursuing academic research careers. 49 of the 81 T32 trainees responded to the survey. We counted the frequency that each barrier or enabler was ranked at a specific level of importance (1, 2, 3, etc). Figure 3 shows the number of times each barrier (upper panel) or enabler (lower panel) was ranked in each position. The size of the square corresponds to the frequency that barrier or enabler was chosen at that rank. The barriers and enablers are listed by decreasing frequency of selection. Cardiovascular T32 trainees identified the primary barriers to pursuing careers in academia as “Perceived Lack of Stability,” “Publish or Perish Competitive Mentality” and “Availability of Positions”. In contrast, they identified “Desire to Contribute to Collective Knowledge,” “Enjoyment of the Spirit of Inquiry,” and “Experience and Skills Gained Through Research” as the primary enablers. In general, respondents responded to a greater number and variety of enablers than barriers.

**Figure 3.**
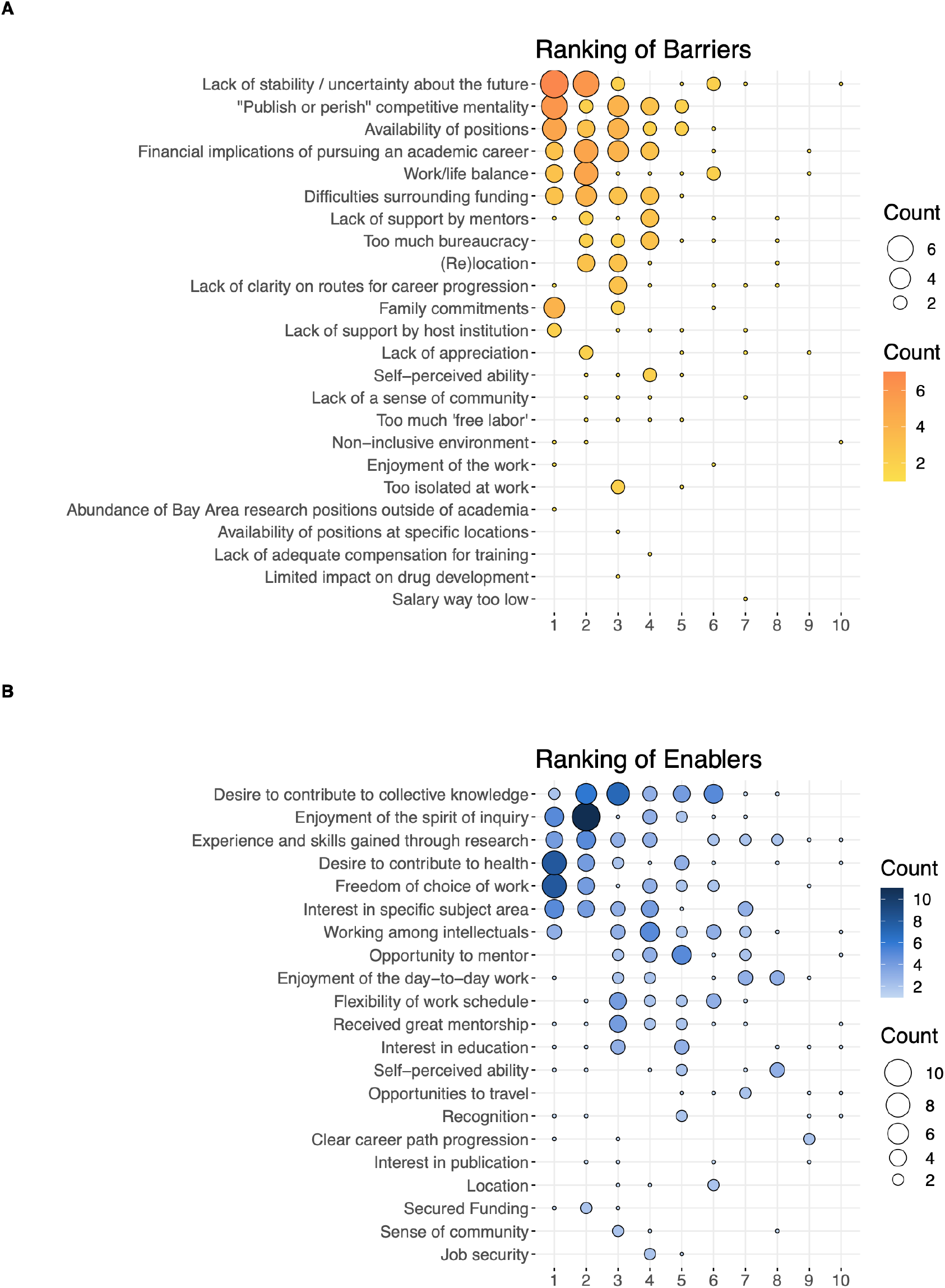
Ranking of barriers and enablers to pursuing careers in academia. Respondents could select any number of barriers or enablers and were then asked to rank them from 1^st^ (1, most significant barriers) to last. Barriers are listed in the upper panel, enablers in the lower panel. The size of the squares in this figure indicates the number of respondents who gave a specific barrier or enabler a specific rank. Data from CV T32 Survey. See accompanying source data “Enablers Rank” and “Barriers Rank” for deidentified data.

## Discussion

Maintaining the pipeline of the next generation of scientists and researchers has proven increasingly difficult. Even at a tier 1 research institution, less than half of the trained postdocs stay in academia, and less than a quarter of trainees attain primarily research faculty positions. Despite postdocs prevalent desire to contribute to collective knowledge and their enjoyment of the spirit of inquiry, they describe being stymied in their pursuit of academic careers by numerous factors including uncertainty about their futures and the cut-throat attitude to success.

One mechanism to enhance retainment in academic careers is to provide the infrastructure offered by programs like T32s. These typically include formal grant writing programs, opportunities to network with visiting professors, guidance on establishing collaborations, public speaking, as well as other professional development opportunities. These opportunities may help reduce the barrier of the perceived lack of stability and make available positions more apparent. In the case at our university, T32 trainees ‘outperformed’ non-T32 trainees, with significantly higher rates of retention in primarily research academic faculty positions. This was also true even amongst our domain-specific cardiovascular T32s, where high numbers of MDs turned down potentially more lucrative roles in industry or as practicing physicians to stay in academia. Our data further suggest that T32 trainees may be particularly valuable in supporting individuals from underrepresented backgrounds. Further studies are needed to further investigate this possibility.

At least in our sample, postdocs voiced strongest concerns about external factors: the uncertainty about finding a position and about their futures and a pervasive sense that their careers hinged on their publication track record. Postdocs were less inhibited about pursuing careers in academia by more personal concerns, such as a sense of community, mentorship, and the feeling that their work was appreciated. This suggests that more should be done to increase postdocs sense of security about their futures – that jobs will be available, and that their careers do not depend solely on generating high-impact publications. For this to be persuasive, hiring committees also need to look beyond applicants’ publication track records to other measures of success such as producing rigorous work, having creative ideas, and being an inclusive mentor, communicator, and educator. It is also worth noting that trainees were more strongly motivated to pursue careers in academia by more ‘lofty’ concepts such as the spirit of inquiry and contributing to collective knowledge, as opposed to, again, more tangible motivators such as the day-to-day work, opportunities to travel, and direct mentorship. This suggests that mentors, funding agencies, and institutions could be doing more to remind trainees of the value of their contributions, and to encourage them to embrace their curiosity and critical analysis.

Although this study provides insight into the motivations and challenges that trainees experience in pursuing academic careers, it is important to acknowledge that our data stems from a limited number of trainees who all experienced the same overall training environment at Stanford University. Additionally, participation in the Barrier and Enabler survey was optional and although 59% of CV trainees responded, their responses may not be fully representative of the population. Another limitation of this study is that our “Non-T32 Postdoc” population may include a small number of T32 trainees. Because of the large size of this population (n = 2,021) and the relatively small proportion of T32 postdocs at Stanford (5%), the impact of T32-trainee presence in this population will be negligible and, if anything, would add noise to our results.

Our results are consistent with the idea that dedicated training programs strengthen the pipeline of young scientists pursuing careers in academic research. In 2020, 0.3% of the NIH budget went towards institutional training grants, yet this study suggests that training programs are a valuable National investment in training motivated and qualified young researchers to pursue scientific discovery for the advancement of human health. However, even trainees in structured and supportive T32 training environments are often inhibited from pursuing academic careers by concerns about the uncertainty of their prospects and also academia’s competitive culture. T32 training programs are a tool to increase the ability, motivation, and diversity of our academic workforce, but institutions also need to prioritize establishing transparent career ladders for faculty positions and look beyond publication metrics in their review of a faculty candidate’s academic merit.

## Materials and Methods

### Study Design and Participants

Data for this study comes from responses to three data sources: 1) data assembled by the institutional Office of Postdoctoral Affairs both from institutional records and research on career outcomes of postdoctoral trainees (OPA Data) who completed their training at Stanford between 2010 and 2020; 2) data from three cardiovascular T32 program training records and progress reports (CV T32 Data), and 3) a survey conducted in 2020 by representatives from the three cardiovascular T32 programs on the barriers and enablers that past and current postdoctoral trainees in the training programs experienced in pursuing careers in academia (CV T32 Survey). This survey was deemed exempt from human subject’s research by the Stanford University Institutional Review Board. See Table 1 for information on the numbers and demographics of individuals in the three datasets (OPA Data, CV T32 Data, CV T32 Survey). Note that non-binary genders are not represented due to current limitations in our institutional process for capturing demographics. Individuals in OPA Data were classified as having confirmed participation in a T32, “T32 Postdocs,” or not, “Non-T32 Postdocs”. For this dataset, only postdocs in relevant health-related departments were included. Note that we were unable to verify the T32 status of all of the postdocs in the “Non-T32 Postdocs” group, so there may still be a small percentage of T32 postdocs in this population (n = 54-108 out of 2,021). Amongst the T32 trainees, the subset who participated in one of our three cardiovascular T32s were also identified “Cardiovascular T32 trainees”. “T32” respondents are individuals who are confirmed to have participated in one of the 15 postdoctoral T32 programs at our institution, including the three cardiovascular T32 programs. “Non-T32 Postdocs” respondents are individuals for whom their T32 participation status is not confirmed. All 81 cardiovascular T32 trainees were asked to complete the survey. Of the 81 cardiovascular T32 trainees for whom data was available, 49 (60%) responded to the CV T32 Survey.

**Table 1.** Demographics of OPA Data and of trainees in three specific cardiovascular T32 programs (CV T32 Program Data.) See accompanying deidentified source data for “OPA Data,” “CV T23 Program Data,” and “CV T32 Survey Data.”

| OPA Data |  |  | CV T32 Program Data | CV T32 Survey Data |
| --- | --- | --- | --- | --- |
| Sex | T32 Postdocs | Non-T32 Postdocs |  |  |
| Female | 126 (50.8%) | 1,049 (51.9%) | 39 (48.1%) | 24 (49%) |
| Male | 122 (49.2%) | 972 (48.1%) | 42 (51.9%) | 25 (51%) |
| **URM Status** |  |  |  |  |
| URM | 32 (12.9%) | 207 (10.2%) |  |  |
| Not URM | 216 (87.1%) | 1,813 (89.7%) |  |  |
| Unknown | 0 (0%) | 1 (0%) |  |  |
| **Total** | 248 (100%) | 2,021(100%) | 81 (100%) | 49 (100%) |

### Measures

In the OPA dataset, the careers of former postdocs were identified using web searches to verify employment via employer websites such as University faculty profiles, research databases, and social media sites such as LinkedIn. Former postdoctoral trainees were identified as pursuing careers in different sectors (Academia, For-Profit, Government, Nonprofit, or Other), as well as whether they were holding faculty or non-faculty positions. They were also identified as pursuing careers that involved “Further Training or Education,” “Primarily Research,” “Primarily Teaching,” “Science/Discipline Related,” or “Not Related to Science/Discipline.” Postdoctoral trainees are considered to be pursuing “Primary Research Faculty” positions if they are identified as faculty, in the sector “Academia”, whose work is “primarily research”. The taxonomy used was a refined version of UCOT 2017^1^ where UCOT 2017 refers to a taxonomy developed by several organizations and described in Silva *et al*, 2019^2^. This taxonomy was developed and applied in order to compare job sectors and career types consistently across institutions. Additionally, individuals from this dataset were identified as either underrepresented minorities or not via Stanford University biodemographic data.

The outcomes of T32 trainees in the three participating programs were identified based on information collected for the programs’ 2020 NIH-mandated progress reports. Data in these progress reports include information from all trainees, including those appointed in previous funding cycles. To collect this information, we either reached out to the trainees directly for information about their current career status or identified their current occupation through online search. Trainees’ current outcomes were categorized as either “Further Training or Education”, “Faculty”, “Industry”, or “Other”. “Other” includes careers in clinical practice, non-higher education teaching, and non-academic research. We also used previous T32 records and progress report data to determine trainee demographic information.

In the CV T32 Survey, trainees were asked to list the barriers and enablers they experienced to pursuing careers in academia from a predefined list of 21 barriers and 21 enablers. These barriers and enablers were identified and pooled from several previous reports on factors that influence early career scientists’ interest in pursuing academic careers.^3–5^ To reduce bias, the order in which the barriers and enablers were listed was randomized. Respondents also had the option to specify up to three additional barriers or enablers. After selecting which barriers or enablers contributed to their decision to pursue a career in academia, respondents were asked to rank the barriers and enablers in order of greatest to least importance. Note that not all cardiovascular T32 trainees participated in the CV T32 Survey. See supplementary information for the full survey.

### Statistical Analysis

Comparisons between different populations of trainees were performed using Chi-Square tests, with p-values < 0.05 considered significant. Bonferroni correction was used to adjust the p-value to correct for multiple comparisons.

## Materials, Methods, and Data Availability

The survey instrument will be included as supplementary material, as will the R Markdown file used to create all three figures. Source data from the OPA Dataset, the CV T32 Data, and the CV T32 Survey are included with this publication and also available on Dryad at https://doi.org/10.5061/dryad.70rxwdc0t. The temporary link for peer review is: https://datadryad.org/stash/share/pFvDz1ZnSBXUsruE9qo_rlyFGSTww86kvLwl4hsX54A.

## Acknowledgments

Corey McCall provided invaluable assistance in identifying the outcomes of postdoctoral trainees in the OPA dataset. We would also like to thank Chantanee Saejao for her efforts in identifying T32 postdocs among the full postdoc population. We are also grateful to Terra Coakley who manages the Research Training in Myocardial Biology at Stanford T32 program and helped collate contact information for several trainees for the survey. Lastly, we would also like to thank Dr. Daniel Bernstein, Dr. Joseph C. Wu, Dr. Thomas Quertermous, Dr. Euan Ashley, Dr. John Pauly, and Dr Koen Nieman, who jointly direct two postdoctoral T32 training programs in cardiovascular research at Stanford University, for their strong support of this study.

## References

1. C.A. Stayart, P. D. Brandt, A. M. Brown, T. Dahl, R. L. Layton, K. A. Petrie, E. N. Flores-Kim, C. G. Peña, C. N. Fuhrmann, G. C. Monsalve, Applying inter-rater reliability to improve consistency in classifying PhD career outcomes. F1000Res. 9;9:8 (2020).

2. E. Silva, A. Mejía, E. Watkins, Where Do Our Graduates Go? A Tool Kit for Tracking Career Outcomes of Biomedical PhD Students and Postdoctoral Scholars. CBE life sciences education. 18. le3. (2019)

3. S. Afonja, D.G. Salmon, S.I. Quailey, W.M. Lambert, Postdocs’ advice on pursuing a research career in academia: A qualitative analysis of free-text survey responses. PLoS ONE 16(5): e0250662. (2021)

4. M. Ålund, N. Emery, B.J.M. Jarrett, et al. Academic ecosystems must evolve to support a sustainable postdoc workforce. Nat Ecol Evol 4, 777–781. (2020)

5. G.S. Gina. Career barriers influencing career success. Career Development International. 21(1):60–84. (2016)

